# A critical residue in the α_1_M2-M3 linker regulating GABA_A_ receptor pore gating by diazepam

**DOI:** 10.1101/2020.10.04.325456

**Authors:** Joseph W. Nors, Shipra Gupta, Marcel P. Goldschen-Ohm

## Abstract

Benzodiazepines (BZDs) are a class of widely prescribed psychotropic drugs. Their anxiolytic and sedative effects are conferred by modulating the activity of GABA_A_ receptors (GABA_A_Rs), which are the primary inhibitory neurotransmitter receptors throughout the central nervous system. However, the physical mechanism by which BZDs exert their effects on the receptor is poorly understood. In particular, BZDs require coapplication with an agonist to effectively open the channel pore, making it difficult to dissect whether the drug has altered either agonist binding or channel gating as these two processes are intimately coupled. To isolate effects on gating we used a spontaneously active gain of function mutant (α_1_L9’Tβ_2_γ_2L_) that is directly gated by BZDs alone in the absence of agonist. In the α_1_L9’T background we explored effects of alanine substitutions throughout the α_1_M2-M3 linker on modulation of the channel pore by the BZD positive modulator diazepam (DZ). The M2-M3 linker is known to be an important element for channel activation. Linker mutations generally impaired unliganded pore opening, indicating that side chain interactions are important for channel gating in the absence of bound agonist. All but one mutation had no effect on the transduction of chemical energy from DZ binding to pore gating. Strikingly, α_1_V279A doubles DZ’s energetic contribution to gating, whereas larger side chains at this site do not. In a wild-type background α_1_V279A enhances DZ-potentiation of currents evoked by saturating GABA, consistent with a direct effect on the pore closed/open equilibrium. Our observations identify an important residue regulating coupling between the BZD site and the pore gate, thereby shedding new light on the molecular mechanism of a frequently prescribed class of psychotropic drugs.

## Introduction

BZDs (e.g. Valium, Xanax) are one of the most widely prescribed psychotropic drugs today. An estimated 100 million adults in the United States fill a prescription for a BZD annually (Agarwal and Landon, 2019; Bachhuber et al., 2016; Olfson et al., 2015). Their anxiolytic and sedative properties are used as therapies for conditions including anxiety, panic, insomnia, seizures, muscle spasms, pain and alcohol withdrawal (Möhler et al., 2002). Although largely effective, BZDs have undesirable effects including tolerance, addiction, dependence, withdrawal symptoms, and are often co-abused with alcohol and opioids (Fluyau et al., 2018; Schmitz, 2016; Jones and McAninch, 2015; Jones et al., 2014). Novel therapies with reduced risks are imperative for safer long-term treatment options.

The therapeutic effects of BZDs are conferred upon binding to and modulating the activity of GABA_A_Rs, which are the primary inhibitory neurotransmitter receptors throughout the central nervous system (Smart and Stephenson, 2019). GABA_A_Rs are part of the superfamily of pentameric ligand-gated ion channels (pLGICs) including homologous glycine (Gly), nicotinic acetylcholine (nACh), serotonin type 3 (5-HT_3_) and zinc-activated receptors as well as prokaryotic homologs (Pless and Sivilotti, 2019; Nemecz et al., 2016; Tasneem et al., 2005; Connolly and Wafford, 2004). Each pentameric GABA_A_R is comprised of subtype-specific combinations of homologous but nonidentical subunits (α_1-6_, β_1-3_, γ_1-3_, δ, ε, π, θ, ρ_1-3_) that together form a central chloride conducting pore (Figure 1) (Olsen and Sieghart, 2008). The most prominent subtype at synapses is comprised of α_1_, β_n_ and γ_2_ subunits (Sieghart and Sperk, 2002). GABA_A_Rs provide a critical balance with excitatory signaling for normal neural function, and not surprisingly their dysfunction is related to disorders such as epilepsy, autism spectrum disorder, intellectual disability, schizophrenia, and neurodevelopmental disorders such as fragile X syndrome (Hernandez and Macdonald, 2019; Gao et al., 2018; Mahdavi et al., 2018; Braat and Kooy, 2015; Vien et al., 2015; Rudolph and Möhler, 2014; Limon et al., 2012; Ramamoorthi and Lin, 2011; Solís-Añez et al., 2007). Although pharmacological manipulation of GABA_A_Rs is a powerful approach to tuning neural signaling, the rational design of novel therapies is challenged by a lack of understanding of the molecular mechanism by which drugs such as BZDs modulate channel behavior.

**Figure 1.**
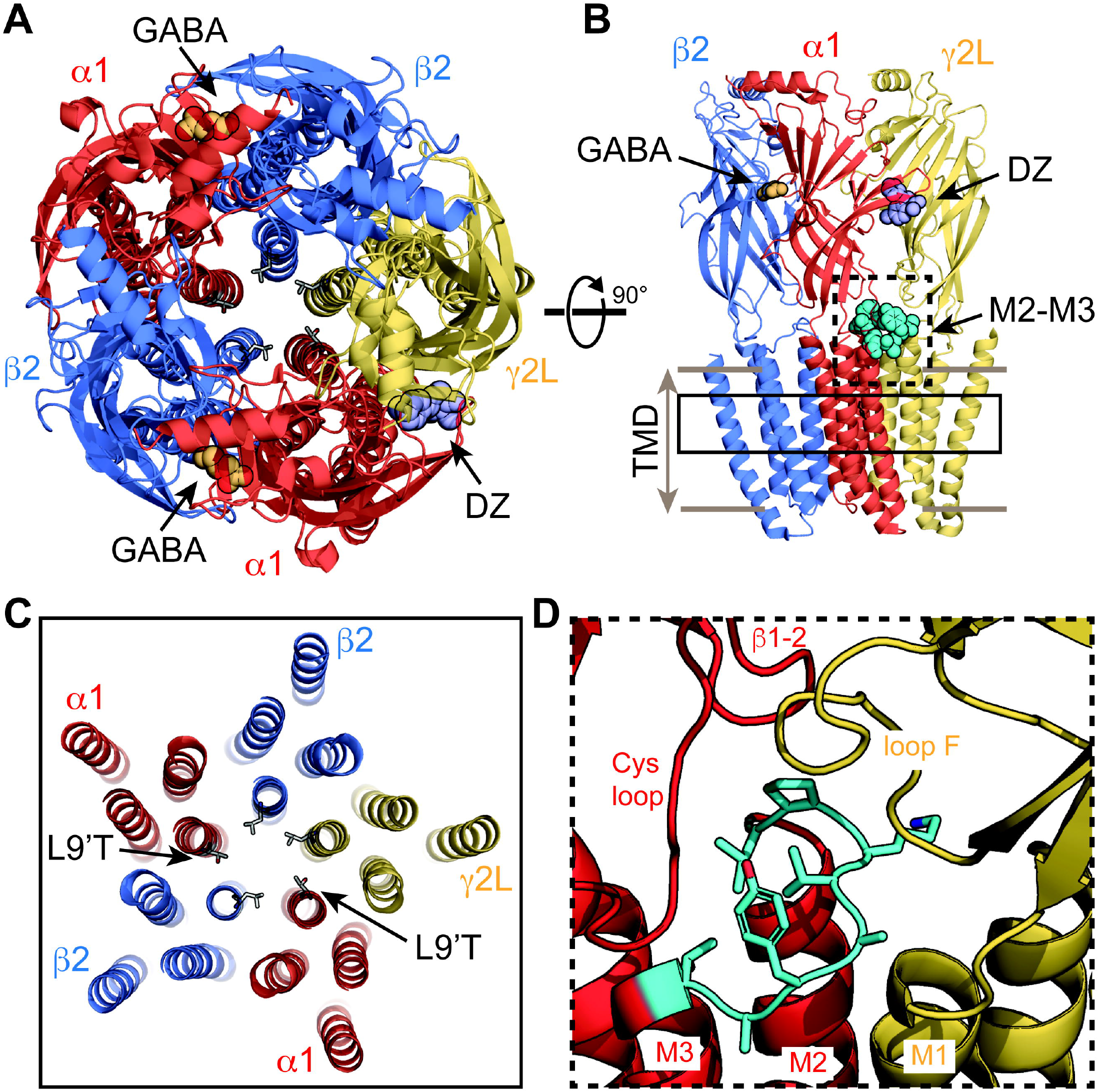
Visual representation of an α_1_β_2_γ_2_ GABA_A_ receptor from cryo-EM map PDB 6×3X. **A-B)** View from the extracellular space (*A*) and parallel to the membrane plane (*B*). Bound GABA and DZ are shown as gold and lavender spheres, respectively. The 9’ pore residue from each subunit is shown in stick representation, all leucine except for the mutation α_1_L9’T which was generated in PyMol as a visualization aid. One of the two α_1_M2-3 linkers is shown as cyan spheres in *B*. **C)** Same view as in *A* for a slice through the transmembrane domains indicated by the solid box in *B*. **D)** Detail for the dashed box in *B*. The α_1_M2-3 linker (L276-T283; rat numbering) is colored cyan with sidechains shown in stick representation.

In conjunction with structural models of mostly homomeric pLGICs, recent cryo-EM models of heteromeric GABA_A_Rs with bound neurotransmitter and BZD provide an important conceptual aid to understand their mechanism of action on these receptors (Kim et al., 2020; Masiulis et al., 2019; Laverty et al., 2019; Phulera et al., 2018; Zhu et al., 2018). Consistent with earlier functional studies (Sigel and Ernst, 2018; Sigel and Lüscher, 2011; Sigel, 2002; Wagner and Czajkowski, 2001; Kucken et al., 2000; McKernan et al., 1998; Pritchett et al., 1989; Gibbs et al., 1985), they show BZDs bound at a specific recognition site in the extracellular domain between α and γ subunits (Figure 1). However, these structures have yet to clarify the mechanism by which BZDs modulate channel activity.

BZDs require coapplication with an agonist to facilitate robust pore opening. This makes it difficult to dissect whether the drug has altered either agonist binding or channel gating as the two processes are intimately coupled (Colquhoun, 1998). This challenge has contributed to differing conclusions postulating that BZDs alter either agonist binding (Goldschen-Ohm et al., 2010; Perrais and Ropert, 1999; Mellor and Randall, 1997; Rogers C J et al., 1994; Vicini et al., 1987), pore gating (Li et al., 2013; Downing et al., 2009; Campo-Soria et al., 2006; Rüsch and Forman, 2005) or an intermediate preactivation step (Goldschen-Ohm et al., 2014; Gielen et al., 2012). The ability of BZDs to potentiate current responses to saturating partial agonists, and also to directly gate gain of function mutants imply that the drug’s effect is not due to changes in binding alone. A combination of effects on both binding and gating as predicted by changes in intermediate gating steps energetically coupled to both binding sites and the pore gate is plausible (Goldschen-Ohm et al., 2014). However, the molecular identity of any such intermediate states remains unclear.

Here we examined the effect of BZDs on pore gating in isolation from any effects on agonist binding using a spontaneously active gain of function mutant (α_1_L9’Tβ_2_γ_2L_) that is directly gated by BZDs alone (Scheller and Forman, 2002; Chang and Weiss, 1999). In the α_1_L9’T background we serially mutated each residue in the α_1_M2-M3 linker to alanine and assessed the effect on modulation of the channel pore by the BZD positive modulator DZ (Valium) in the absence of agonist. The M2-M3 linker is a loop following the pore lining M2 helix that together with several other important interfacial loops defines the region connecting extracellular ligand binding and transmembrane domains known to be crucially involved in the gating process of pLGICs (Figure 1). Structural models show that agonist binding is associated with an outward displacement of the M2-M3 linker from the central pore axis (Nemecz et al., 2016). Mutations in the linker generally impair gating by agonists (Kash et al., 2003; O’Shea and Harrison, 2000), and some are associated with genetic diseases such as epilepsy in GABA_A_Rs (Hales et al., 2006; Hernandez and Macdonald, 2019) or hyperekplexia in GlyRs (Bode and Lynch, 2014). However, the role of the M2-M3 linker in drug modulation by BZDs is less understood.

We show that alanine substitutions throughout the α_1_M2-M3 linker generally impair unliganded pore gating, whereas they have little effect on the efficiency by which chemical energy from DZ binding is transmitted to the pore gate. The notable exception is α_1_V279A which more than doubles DZ’s energetic contribution to pore opening in both gain of function and wild-type backgrounds, possibly by removal of a steric hindrance. The effect of this mutation is largely explained by a simple model postulating direct effects on the pore gating equilibrium. Our observations identify an important residue in the α_1_M2-M3 linker regulating the efficiency of BZD-modulation in GABA_A_Rs.

## Results

### Alanine substitutions in the α_1_M2-M3 linker generally impair unliganded gating

In the α_1_L9’Tβ_2_γ_2L_ (α_1_L9’T or L9’T) gain of function background we assessed the effects of alanine substitutions in the α_1_M2-M3 linker on unliganded and GABA-evoked channel activity. Briefly, Xenopus laevis oocytes were co-injected with mRNA for α, β and γ subunits in a 1:1:10 ratio, and current responses to microfluidic application of ligands was recorded in two-electrode voltage clamp. Each recording consisted of a series of 10 second pulses of GABA at various concentrations bookended by 10 second pulses of 1 mM picrotoxin (PTX) (Figure 2). Current block by the pore blocker PTX was used to assess the amount of spontaneous activity and to correct for drift or rundown over the course of the experiment (see Supplementary Figure 2-1). Normalized concentration-response relations for the additional GABA-evoked current above the spontaneous current baseline are shown in Figure 2. As compared to α_1_L9’T, all of the alanine mutations exhibited an increase in the EC_50_ and/or steepness of the GABA concentration-response relation (Figure 2; Supplementary Figure 2-2). The increased EC_50_ is generally consistent with previous reports of α_1_M2-M3 linker mutations in a wild-type background (O’Shea and Harrison, 2000) and imply that side chain interactions in this region are important for channel gating by agonists. The steepness of the concentration-response relation for α_1_L9’T was fairly shallow, with a hill slope slightly below one. This could reflect an increased efficiency of mono-liganded gating in the gain of function background. In such a case, the steeper hill slopes conferred by α_1_L276A, α_1_V279A and α_1_Y281A may be due to a reduction in the efficiency of mono-vs di-liganded gating, although this was not verified.

**Figure 2.**
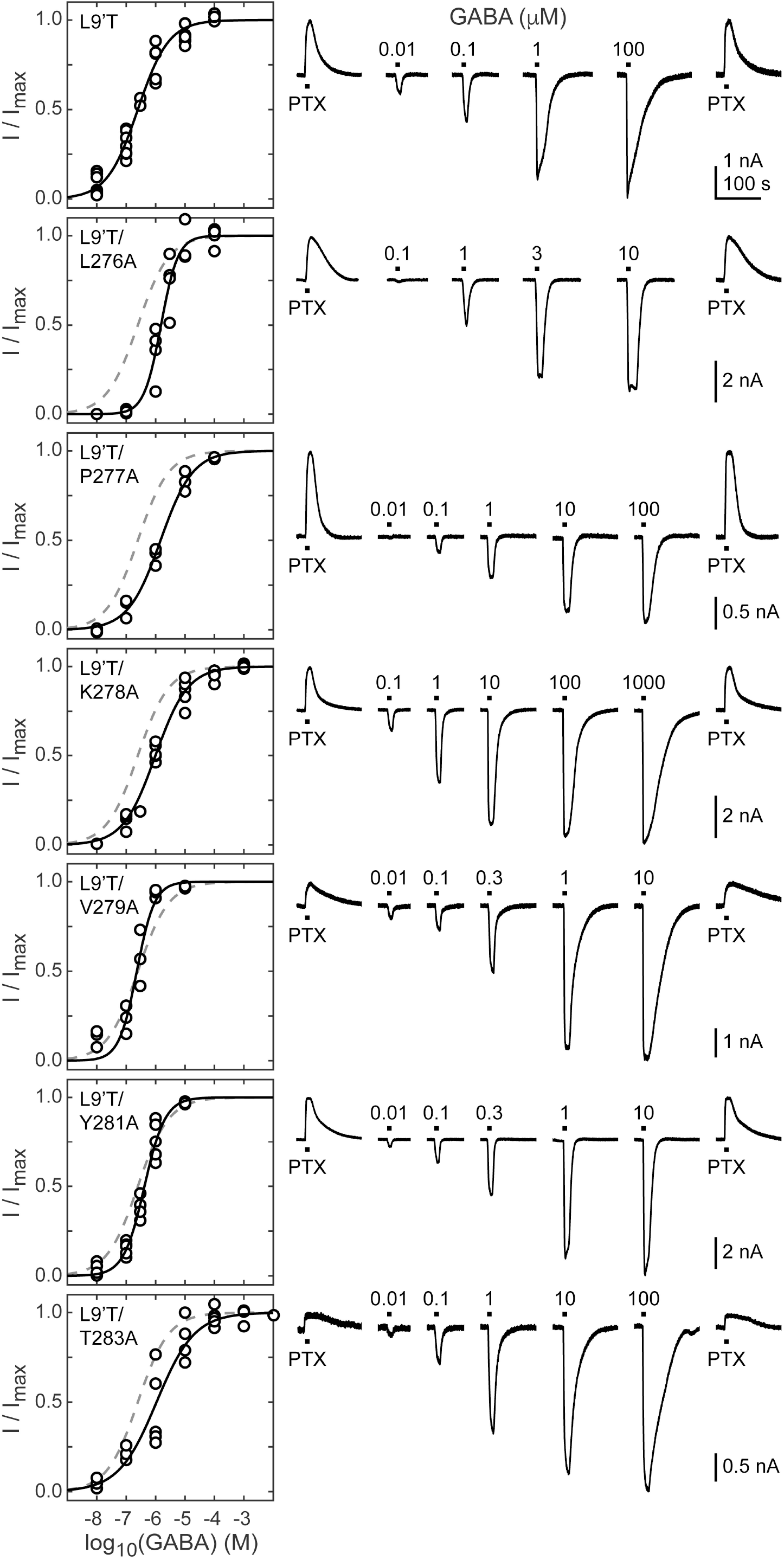
Spontaneous PTX-sensitive and GABA-evoked currents for α_1_M2-M3 linker alanine substitutions in the gain of function α_1_L9’Tβ_2_γ_2L_ background. ***left*)** Summary of normalized GABA concentration-response relations for GABA-evoked currents with the zero current baseline set to the level of spontaneous activity. Solid line is a fit of the combined data across oocytes to Equation 1, and the dashed line is the fit for the L9’T background. Fit parameters are EC_50_, hill slope (# oocytes): L9’T = 0.25 μM, 0.83 (7); L9’T/L276A = 1.53 μM, 1.51 (4); L9’T/P277A = 1.48 μM, 0.80 (3); L9’T/K278A = 0.99 μM, 0.79 (5); L9’T/V279A = 0.23 μM, 1.44 (4); L9’T/Y281A = 0.42 μM, 1.23 (5); L9’T/T283A = 1.08 μM, 0.67 (5). Parameters for fits to individual oocytes are summarized in Supplementary Figure 2-2. ***right*)** Example currents in response to 10 second pulses of the pore blocker PTX (1 mM) or GABA (concentration in micromolar indicated above each pulse). Responses to GABA were bookended by application of PTX to assess the amount of spontaneous current and to normalize any drift or rundown during the experiment (see Supplementary Figure 2-1).

The level of unliganded channel activity was determined from the ratio of the spontaneous current amplitude as assessed by block with PTX (I_PTX_) to the maximal current amplitude evoked with saturating GABA (I_GABA-max_) (Figure 3A). The unliganded open probability was estimated as the product of the ratio I_PTX_/I_GABA-max_ and the open probability in saturating GABA (P_o-GABA-max_). Under the assumption that P_o-GABA-max_ is similar for all constructs, all α_1_M2-M3 linker mutations except for α_1_P277A reduced the unliganded open probability by approximately two-fold (Figure 3B). P_o-GABA-max_ is ∼0.85 in wild-type channels (Keramidas and Harrison, 2010), and likely to be closer to 1.0 in a gain of function background such as α_1_L9’T that is further known to desensitize much more slowly than wild-type (Scheller and Forman, 2002). Thus, any reduction in P_o-GABA-max_ for the linker mutations should only further reduce their estimated unliganded open probability. In summary, these data show that alanine mutations in the α_1_M2-M3 linker generally impair channel gating by increasing GABA EC_50_ and/or reducing unliganded pore opening. This suggests that α_1_M2-M3 linker side chain interactions play an important role in the closed vs. open pore equilibrium even in the absence of agonist.

**Figure 3.**
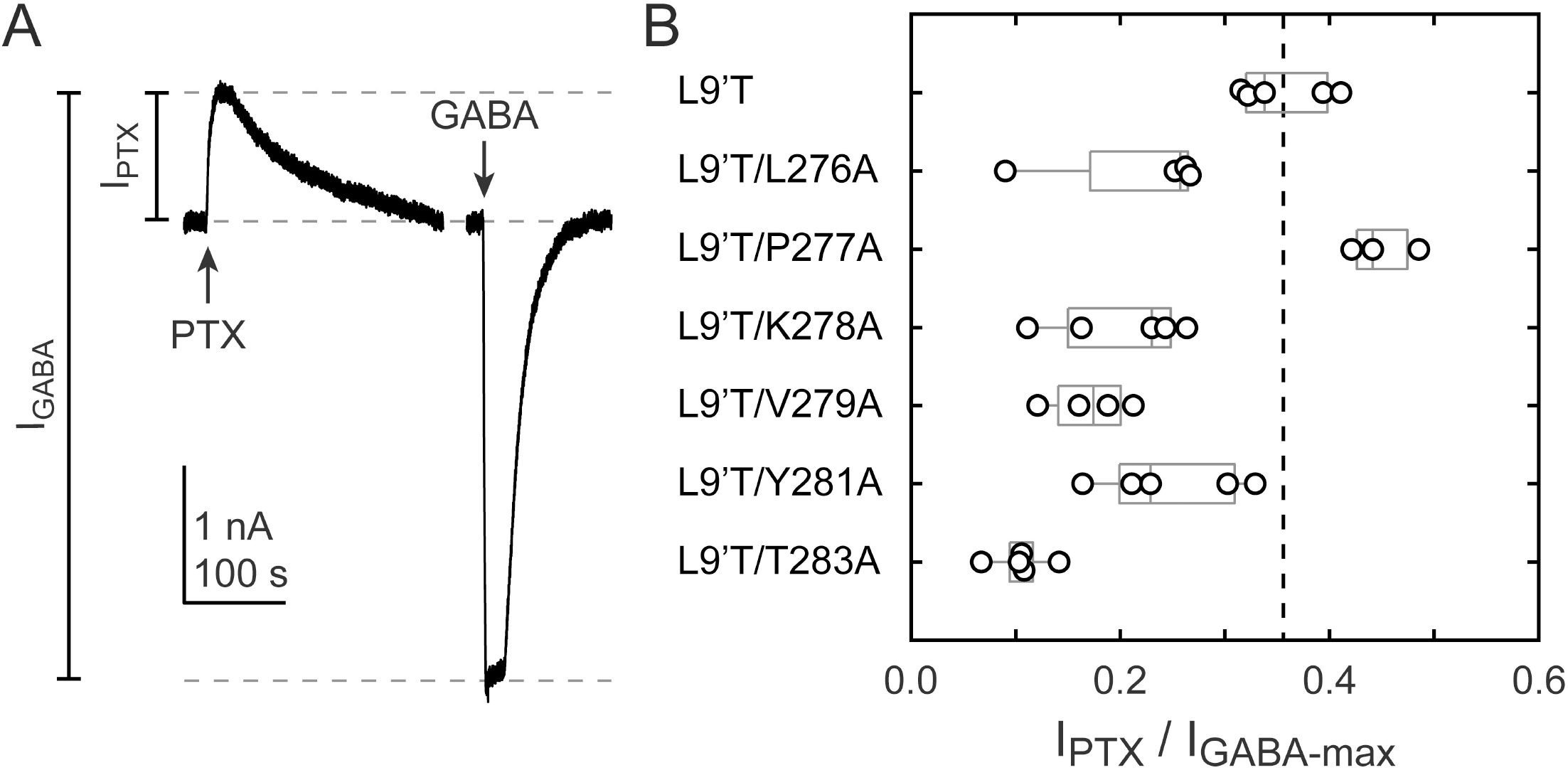
Ratio of PTX-sensitive to maximal GABA-evoked current amplitude. **A)** Example currents from α_1_L9’T/V279Aβ_2_γ_2L_ receptors elicited by 10 second pulses of either 1 mM PTX or 10 μM GABA. **B)** Summary of the ratio of PTX-sensitive to maximal GABA-evoked current amplitude for individual oocytes. Gray box plots indicate the median and 25^th^ and 75^th^ percentiles. The vertical dashed line is the mean for L9’T.

### All but one alanine substitution in the α_1_M2-M3 linker have little effect on activation by DZ relative to unliganded activity

BZD positive modulators alone evoke channel opening with extremely low probability (Campo-Soria et al., 2006), thus necessitating coapplication with an agonist to obtain robust currents from wild-type receptors. However, dissecting the effects of BZDs on either agonist binding or channel gating is severely challenged in the presence of an agonist because the two processes are intimately coupled (Colquhoun, 1998). To isolate effects on gating apart from any effects on agonist binding we examined the effect of alanine substitutions in the α_1_M2-M3 linker on DZ-evoked currents in the background of the gain of function mutation α_1_L9’T in the absence of agonist. Concentration-response relationships for currents evoked by 10 second pulses of DZ were bookended by 10 second pulses of 1 mM PTX to assess spontaneous activity (Figure 4). Drift or rundown was corrected as described for GABA-evoked currents (Supplementary Figure 2-1). Normalized concentration-response relations for the additional DZ-elicited current above the spontaneous current baseline were fit to Equation 1. Alanine substitutions in the α_1_M2-M3 linker have no obvious effect on DZ concentration-response relations with the exception of α_1_L276A which confers a right-shift and reduction in steepness (Figure 4; Supplementary Figure 4-1). Reports for the EC_50_ of DZ potentiation in wild-type receptors are similar to the DZ EC_50_ for α_1_L9’T receptors reported here and in previous studies (Li et al., 2013; Campo-Soria et al., 2006; Rüsch and Forman, 2005; Walters et al., 2000). This is consistent with the idea that both DZ-modulation of GABA-evoked currents and direct gating of the L9’T mutant reflect DZ binding to the same site. The ratio of the maximal DZ-evoked current amplitude (I_DZ-max_) to PTX-sensitive current amplitude (I_PTX_) is a measure for how well DZ activates the channel relative to its spontaneous unliganded activity. This ratio is largely independent of alanine substitution in the α_1_M2-M3 linker with the notable exception of α_1_V279A (Figure 5). Thus, apart from α_1_V279A, DZ-evoked currents are predictably proportional to the amount of unliganded activity. This suggests that α_1_V279 has a unique role in DZ-modulation, whereas side chains of other linker residues are involved little or not at all.

**Figure 4.**
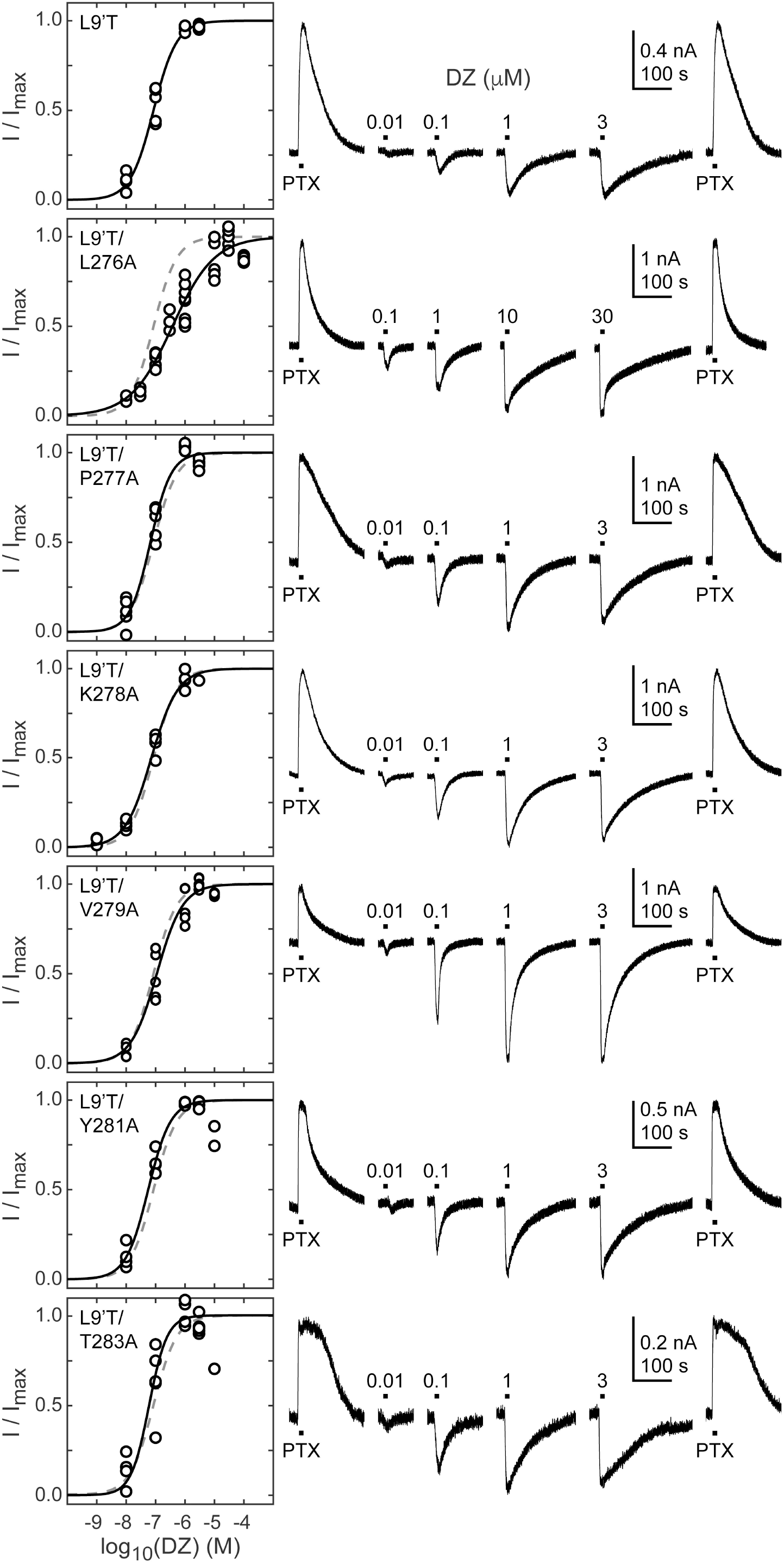
Spontaneous PTX-sensitive and DZ-evoked currents for α_1_M2-M3 linker alanine substitutions in the gain of function α_1_L9’Tβ_2_γ_2L_ background. ***left*)** Summary of normalized DZ concentration-response relations for DZ-evoked currents with the zero current baseline set to the level of spontaneous activity. Solid line is a fit of the combined data across oocytes to Equation 1, and the dashed line is the fit for the L9’T background. Reduced peak responses to 10 μM DZ for L9’T/Y281A and L9’T/T283A were omitted from the fits. Fit parameters are EC_50_, hill slope (# oocytes): L9’T = 85 nM, 1.10 (5); L9’T/L276A = 380 nM, 0.61 (8); L9’T/P277A = 67 nM, 1.24 (5); L9’T/K278A = 72 nM, 0.97 (4); L9’T/V279A = 110 nM, 0.98 (5); L9’T/Y281A = 56 nM, 1.14 (4); L9’T/T283A = 60 nM, 1.42 (6). Parameters for fits to individual oocytes are summarized in Supplementary Figure 4-1. ***right*)** Example currents in response to 10 second pulses of the pore blocker PTX (1 mM) or DZ (concentration in micromolar indicated above each pulse). Responses to DZ were bookended by application of PTX to assess the amount of spontaneous current and to normalize any drift or rundown during the experiment (see Supplementary Figure 2-1).

**Figure 5.**
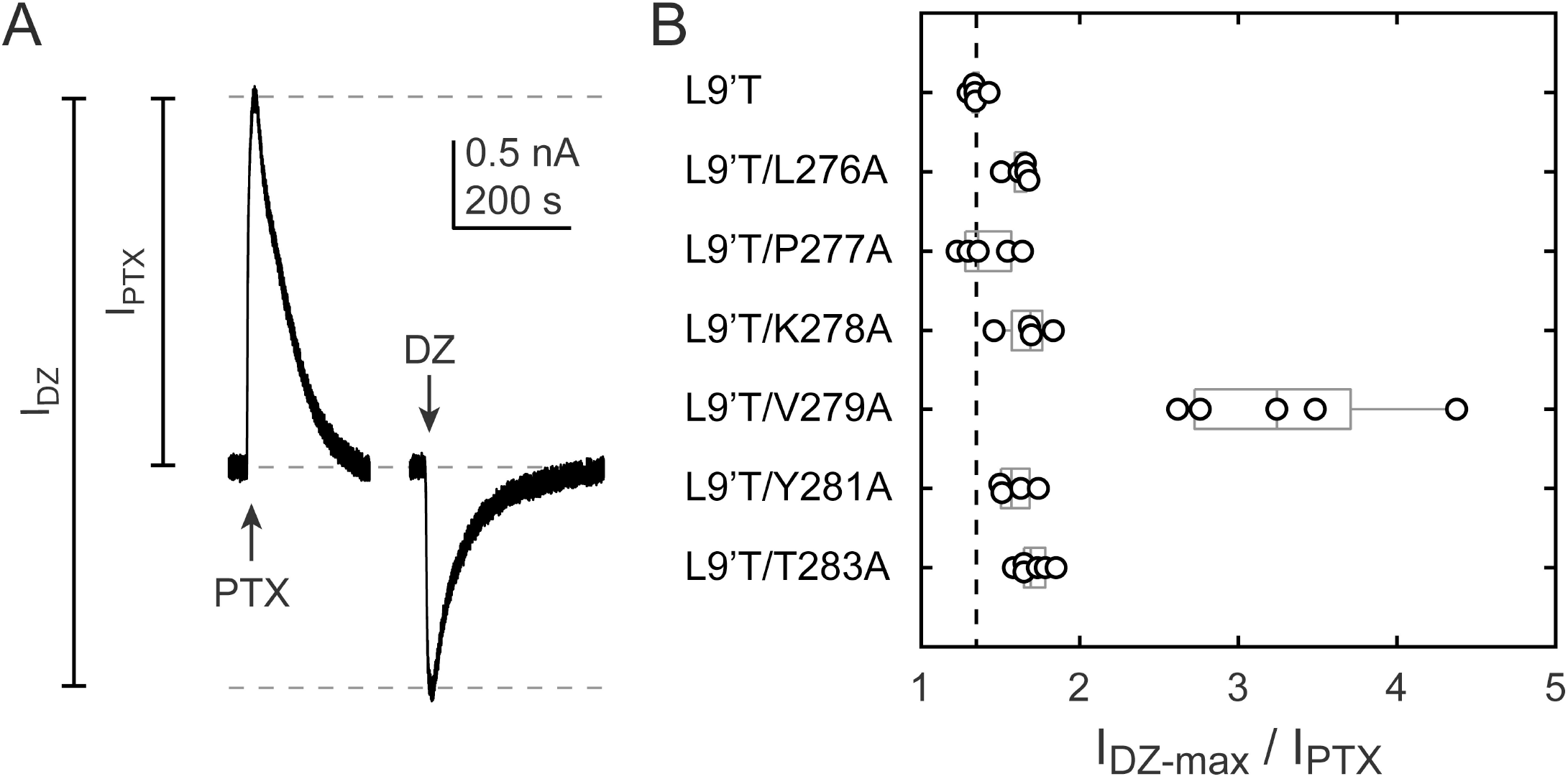
Ratio of maximal DZ-evoked to PTX-sensitive current amplitude. **A)** Example currents from α_1_L9’T/K278Aβ_2_γ_2L_ receptors elicited by 10 second pulses of either 1 mM PTX or 1 μM DZ. **B)** Summary of the ratio of maximal DZ-evoked to PTX-sensitive current amplitude for individual oocytes. Gray box plots indicate the median and 25^th^ and 75^th^ percentiles. The vertical dashed line is the mean for L9’T.

### Dependence of DZ-gating on charge and volume at α_1_V279

The M2-M3 linker has both high sequence and structural similarity in pLGICs. The valine at position 279 in the GABA_A_R α_1_ subunit is located near the center of the linker and is nearly always an aliphatic residue in different GABA_A_R subunits as well as in subunits of other pLGICs, with the exception of AChR, where it is a threonine. Even where sequences differ, structural models of the M2-M3 linker for multiple pLGICs are highly similar. To explore the side chain properties relevant to DZ-gating, we examined the effect of introducing a charged aspartate or bulky tryptophan at position 279 in the α_1_L9’T background. We recorded unliganded PTX-sensitive and GABA- or DZ-evoked currents as described above and compared their relative current amplitudes and concentration-response relations (Figure 6). The negative charge introduced by α_1_V279D severely inhibits unliganded gating, exhibiting very little PTX-sensitive current in relation to robust currents evoked with saturating GABA. In contrast, addition of the bulky side chain α_1_V279W only slightly impairs unliganded gating to a similar degree as substitution with alanine. These data suggest that side chain volume at this position is less critical for unliganded gating, whereas introduction of a negative charge nearly abolishes spontaneous activity despite the α_1_L9’T pore mutation. Consistent with impaired pore gating, α_1_V279D increases the GABA EC_50_ by ∼100-fold.

**Figure 6.**
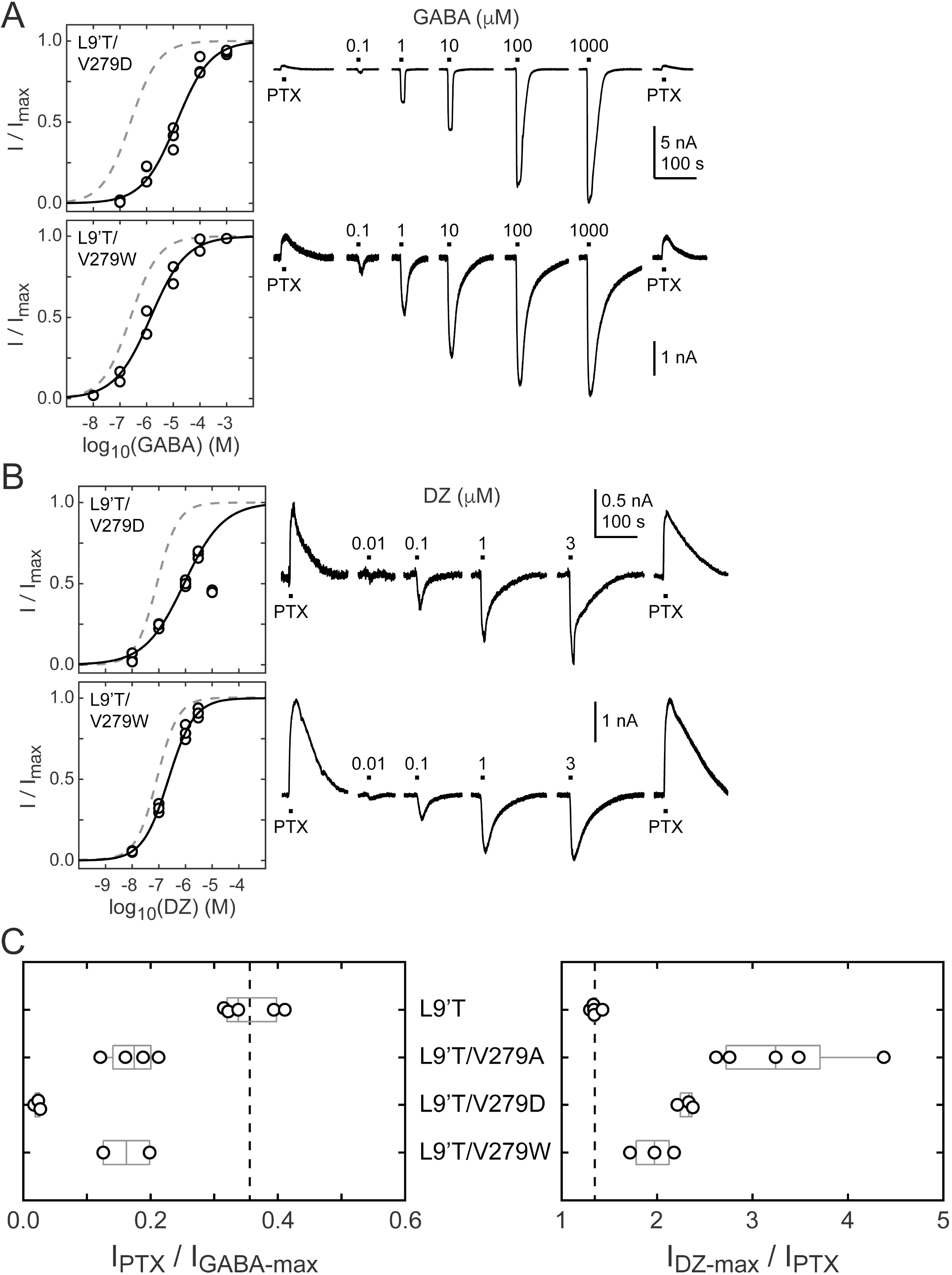
Spontaneous PTX-sensitive and GABA- or DZ-evoked currents for α_1_V279D and α_1_V279W in the gain of function α_1_L9’Tβ_2_γ_2L_ background. **A)** Summary of normalized GABA concentration-response relations for GABA-evoked currents with the zero current baseline set to the level of spontaneous activity (*left*). Solid line is a fit of the combined data across oocytes to Equation 1, and the dashed line is the fit for the L9’T background. Fit parameters are EC_50_, hill slope (# oocytes): L9’T/V279D = 13 μM, 0.68 (3); L9’T/V279W = 1.4 μM, 0.65 (2). Example currents in response to 10 second pulses of the pore blocker PTX (1 mM) or GABA (concentration in micromolar indicated above each pulse) (*right*). **B)** Same as in *A* except for DZ-evoked currents. Responses from L9’T/V279D to 10 μM DZ were excluded from the fit. Fit parameters are EC_50_, hill slope (# oocytes): L9’T/V279D = 0.88 μM, 0.58 (3); L9’T/V279W = 0.23 μM, 0.90 (3). **C)** Summary of the ratios of PTX-sensitive and either maximal GABA- or DZ-evoked current amplitudes for individual oocytes. Gray box plots indicate the median and 25^th^ and 75^th^ percentiles. The vertical dashed line is the mean for L9’T.

The ratio of DZ-evoked to PTX-sensitive current amplitude is only slightly increased by α_1_V279W, whereas it is further increased by α_1_V279D, albeit not to the extent of α_1_V279A (Figure 6C). However, we were unable to reach saturation for DZ-evoked currents from α_1_V279D due to a reduction in peak current amplitude at higher DZ concentrations consistent with occupation of a secondary lower affinity inhibitory site (Figure 6B). Thus, our observations for α_1_V279D reflect a lower limit on channel activity evoked by DZ binding. Even if α_1_V279D has an appreciable effect on the ratio of DZ-evoked to unliganded current, the overall effect of this mutation on channel function is likely to be dominated by its inhibition of pore gating. In contrast, α_1_V279W has similar effects to alanine substitutions at other linker residues. These data suggest that DZ-gating is specifically enhanced by a reduction of side chain volume at position 279.

### The mutation α_1_V279A increases DZ’s energetic contribution to pore gating

To estimate the amount of chemical energy from DZ binding that is transmitted to the pore gate, we employed a simple model of channel gating between closed (C) and open (O) pore states in both unliganded and DZ-bound conditions (Figure 7A). The model assumes that gating can be approximated with a single closed and open state in each liganded condition. The free energy differences for pore gating in unliganded (ΔG_0_) and DZ-bound (ΔG_1_) conditions were calculated according to Equations 2-3 under the assumption that P_open-GABA-max_ ∼ 1. A comparison of ΔG_0_ versus ΔG_1_ shows that DZ binding confers a uniform ΔΔG_DZ_ = ΔG_1_ - ΔG_0_ = 0.7 kT to the pore gating equilibrium independent of α_1_M2-M3 linker mutation with the notable exception of α_1_V279A which more than doubles ΔΔG_DZ_ (Figure 7B,C). Importantly, this result is largely independent of our assumption for P_open-GABA-max_ (see bold vs. light gray symbols in Figure 7B). These observations suggest that coupling between the BZD site and pore gate is relatively independent of individual α_1_M2-M3 linker side chain interactions except for α_1_V279, which natively hinders DZ-gating as compared to its substitution with alanine. Strikingly, introduction of a bulky tryptophan or charged aspartate at position 279 have much smaller effects on ΔΔG_DZ_, consistent with the idea that the small side chain volume of alanine is the most relevant factor for increased DZ efficiency.

**Figure 7.**
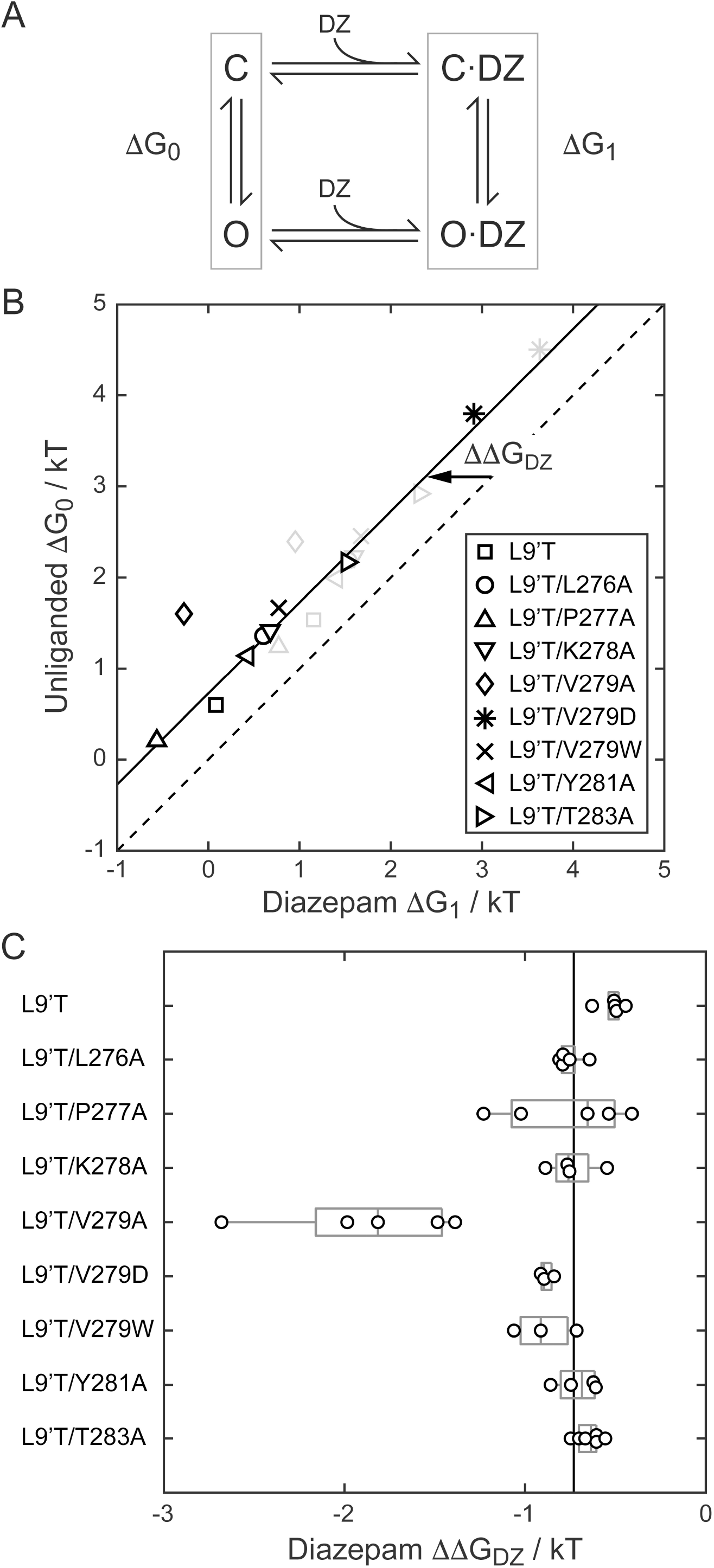
A critical residue in the α_1_M2-M3 linker regulating DZ’s energetic contribution to pore gating. **A)** A simple model approximating channel gating between closed (C) and open (O) pore states in both unliganded and DZ-bound conditions. **B)** Relationship between gating free energy in unliganded (ΔG_0_) and DZ-bound (ΔG_1_) conditions for α_1_L9’Tβ_2_γ_2L_ receptors and α_1_M2-M3 linker mutations assuming *P*_*o*−*GABA*−*max*_ = 1 (bold symbols) or *P*_*o*−*GABA*−*max*_ = 0.5 (light gray symbols) (Equations 2-3). Points are the mean across oocytes. The dashed line of symmetry reflects ΔG_0_ = ΔG_1_ where DZ would have no effect on pore gating. The solid line is a fit to Δ*G*_1_ = Δ*G*_0_ + ΔΔ*G*_*DZ*_ for all of the data points except the outlier α_1_V279A given *P*_*open*−*max*_ = 1. The good description of the data suggests that DZ’s energetic contribution to pore gating is the same for all of the constructs on this line, estimated as ΔΔ*G*_*DZ*_ = 0.7 *kT*. In contrast, α_1_V279A more than doubles the energy that DZ binding transmits to the pore gate. A comparison of the bold and light gray symbols shows that reducing *P*_*o*−*GABA*−*max*_ to 0.5 primarily shifts the data points along the fitted line with only minor changes in ΔΔ*G*_*DZ*_, indicating that our assumption for the value of *P*_*o*−*GABA*−*max*_ is not critical for interpretation of ΔΔ*G*_*DZ*_. **C)** Summary of ΔΔ*G*_*DZ*_ for individual oocytes. Gray box plots indicate the median and 25^th^ and 75^th^ percentiles. The vertical line is the position of linear fit in *B* corresponding to 0.7 kT.

### The mutation α_1_V279A enhances DZ-potentiation of GABA-evoked currents in a wild-type background

If gating in the α_1_L9’T background is relevant to neurotransmitter-driven gating in native receptors, then the mutation α_1_V279A should both impair channel opening and enhance DZ-potentiation of GABA-evoked currents in a wild-type background. To test this, we compared GABA-evoked current responses in α_1_β_2_γ_2L_ (wild-type) and α_1_V279Aβ_2_γ_2L_ (V279A) receptors, as well as the ability of 1 μM DZ to potentiate these responses. First, GABA EC_50_ is increased by α_1_V279A, consistent with impaired unliganded gating (Figure 8A). This effect is similar, although slightly larger than previously observed in α_2_β_1_γ_2S_ receptors (O’Shea and Harrison, 2000). Second, DZ potentiates GABA-evoked peak current responses to a greater extent in V279A than in wild-type receptors, consistent with enhanced coupling between the BZD site and pore gate in the mutant (Figure 8B,C). In contrast to wild-type, DZ also potentiates V279A currents evoked with a saturating concentration of GABA. Together, these data suggest that the additional DZ-modulation conferred by α_1_V279A is both independent of agonist association and additive to DZ-potentiation in wild-type receptors.

**Figure 8.**
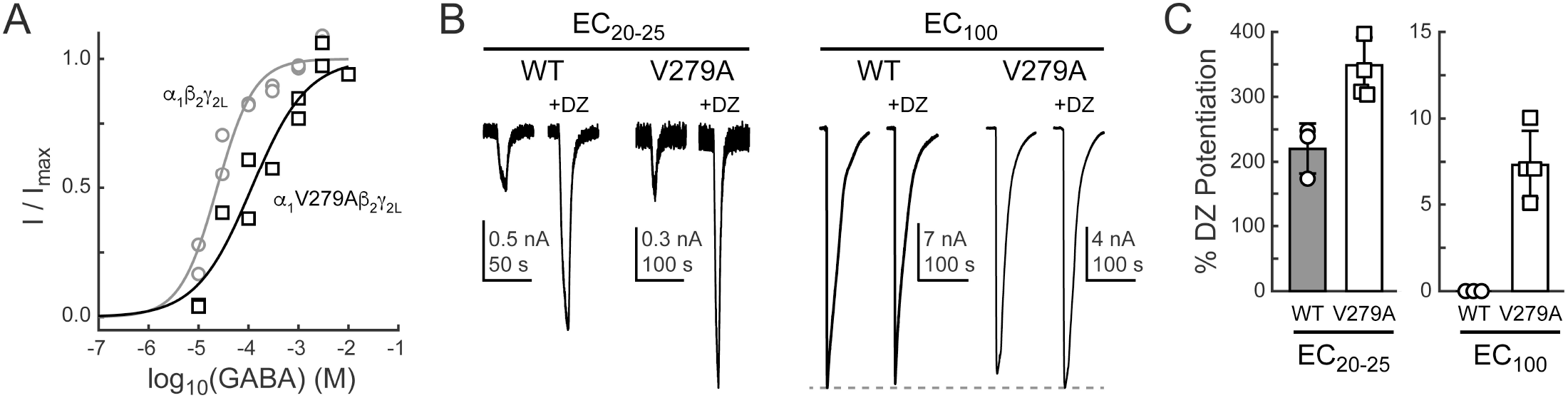
The mutation α_1_V279A enhances DZ-potentiation of GABA-evoked current amplitudes in a wild-type (WT) α_1_β_2_γ_2L_ background. **A)** Normalized GABA concentration-response relations for WT (circles, 2 oocytes) and V279A (squares, 2 oocytes). Solid lines are fits to Equation 1 for all oocytes combined. Fit parameters are EC_50_, hill slope (# oocytes): WT = 24 μM, 1.2 (2); V279A = 116 μM, 0.8 (2). **B)** Potentiation of current amplitudes evoked by 10 second pulses of either subsaturating EC_20-25_ (WT: 10 μM, V279A: 30 μM) or saturating EC_100_ (WT: 3 mM, V279A: 3-10 mM) GABA by 1 μM DZ. **C)** Summary of potentiation as shown in *B* for individual oocytes.

### α_1_L9’T and α_1_V279A have independent and additive effects on the pore gating equilibrium

To further explore the idea that α_1_V279A confers its effects primarily by altering pore gating in DZ-bound receptors, we asked whether a simple Monod-Wyman-Changeux (MWC) model of receptor behavior can account for our observations (Figure 9A). This model has been widely used to describe pseudo steady-state behavior in GABA_A_ receptors (Steinbach and Akk, 2019; Rüsch and Forman, 2005; Chang and Weiss, 1999). For comparison with model simulations we estimated open probability (P_o_) as a function of GABA or DZ concentration (black curves in Figure 9B) from fits of Equation 1 to the data shown in Figures 2, 4 and 8 where the minimum and maximum P_o_ were scaled according to our observed PTX-sensitive and GABA- or DZ-evoked current amplitudes. In the α_1_L9’T background the maximum GABA-evoked P_o_ was allowed to vary during optimization, and the minimum unliganded P_o_ was scaled relative to the maximum by the observed mean ratio of I_PTX_ / I_GABA-max_ (Figure 3). The maximum DZ-evoked P_o_ was scaled relative to the minimum unliganded P_o_ by the observed mean ratio of I_DZ-max_ / I_PTX_ (Figure 5). The maximal P_o_ was set to 0.85 for α_1_β_2_γ_2L_ receptors (Keramidas and Harrison, 2010) and allowed to vary for α_1_V279Aβ_2_γ_2L_ receptors.

**Figure 9.**
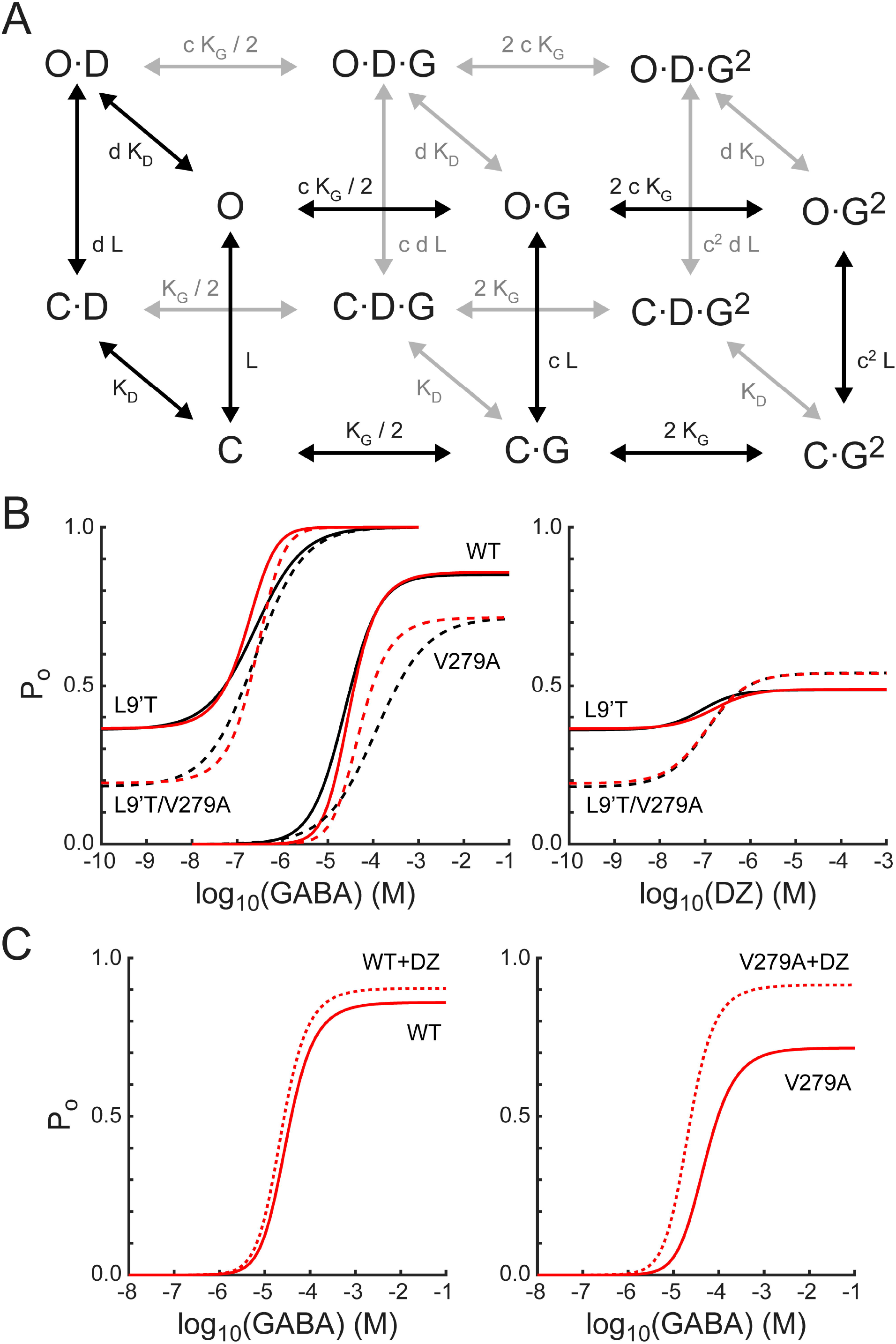
A simple MWC model of channel gating largely accounts for the observed effects of α_1_L9’T and α_1_V279A via independent and additive effects on the pore gating equilibrium. **A)** The model depicts channel gating between closed (C) and open (O) states with independent binding of two GABA (G) and one DZ (D) molecule. L is the ratio of closed to open state probabilities in the absence of ligand, K_G_ and K_D_ are the respective dissociation constants for GABA or DZ, and c and d are the respective factors by which GABA or DZ binding influence channel opening. The probability to be in an open state is given by Equation 4. **B)** Estimated open probability (P_o_) from the data in Figures 2-5 (black) overlaid with model simulations (Equation 4, red). See main text for a detailed description. Model parameters are: L_L9’T_ = 1.8; L_L9’T/V279A_ = 4.2; L_WT_ = 18000; K_G_ = 57 μM; K_D_ = 200 nM; c = 0.0031; d = 0.60; d_V279A_ = 0.20. **C)** The model’s prediction for potentiation of WT and V279A GABA-evoked responses by 1 μM DZ.

The model assumes independent GABA and DZ binding. Model parameters were optimized by globally minimizing the sum of squared residuals between simulated (Equation 4) and estimated P_o_ as a function of GABA or DZ concentration for α_1_L9’Tβ_2_γ_2L_, α_1_L9’T/V279Aβ_2_γ_2L_ and α_1_β_2_γ_2L_ receptors (Figure 9B). Responses to α_1_V279Aβ_2_γ_2L_ receptors were not considered during optimization and treated as a model prediction. The mutations α_1_L9’T and α_1_V279A were allowed to perturb the parameter L with the assumption that their effects are independent in the L9’T/V279A double mutant. For wild-type receptors L was fixed according to its approximate relation with EC_50_ as *L*_*WT*_ = *L*_*L*9*T*_ (*EC*_50,*WT*_ /*EC*_50,*L*9*T*_)^2^ (Steinbach and Akk, 2019; Akk et al., 2018; Scheller and Forman, 2002; Chang and Weiss, 1999). The value of d was assumed to be the same for L9’T and wild-type receptors and allowed to vary in the presence of the α_1_V279A mutation. The resulting optimized model parameters (Figure 9 caption) were highly constrained by the data (i.e. estimated P_o_), and the model does a fairly good job of describing our observed responses to either GABA or DZ, differing somewhat in the steepness of their concentration dependence (Figure 9B). Furthermore, the model qualitatively accounts for observations from α_1_V279Aβ_2_γ_2L_ receptors even though they were not considered during optimization. The model’s predicted maximum P_o_ approached 1.0 in the α_1_L9’T background, consistent with a gain of function phenotype. In contrast, maximal P_o_ was reduced in α_1_V279Aβ_2_γ_2L_ as compared to wild-type receptors, as expected for a reduction in unliganded activity. These results suggest that α_1_L9’T and α_1_V279A have independent and additive effects on the gating equilibrium, and that for α_1_V279A this effect depends on DZ occupancy.

Although responses to GABA or DZ alone are largely explained by this model, DZ-potentiation of GABA-evoked responses in a wild-type background are either underestimated or overestimated for subsaturating or saturating GABA concentrations, respectively (Figure 9C). Previous applications of this model to BZD-modulation in gain of function backgrounds predict a larger left-shift of the P_o_ curve in the presence of drug more similar to that observed, in part due to a smaller value of d (Campo-Soria et al., 2006; Downing et al., 2009; Rüsch and Forman, 2005). However, reducing d also enhances the predicted potentiation of responses to saturating GABA, contrary to that observed for wild-type receptors. Thus, the model is either too simplistic or its assumptions too strict to describe all aspects of DZ-potentiation in wild-type channels that rapidly desensitize, and which may involve drug modulation of agonist binding and/or intermediate gating steps (Goldschen-Ohm et al., 2014; Gielen et al., 2012). Nonetheless, the model does predict an increase in DZ-potentiation of responses evoked by both subsaturating and saturating GABA for α_1_V279A, qualitatively similar to that observed.

## Discussion

The main conclusions of this study are as follows: First, in the α_1_L9’T gain of function background alanine substitutions in the α_1_M2-M3 linker generally impair unliganded pore opening, indicating that side chain interactions with the linker are important for gating even in the absence of bound agonist. Second, the same mutations have no effect on the amount of chemical energy from DZ binding transmitted to the pore gate, except for α_1_V279A which more than doubles DZ’s energetic contribution to pore gating. Thus, α_1_V279 plays a crucial role to natively hinder drug modulation as compared to its substitution with alanine. Third, introduction of a bulky tryptophan or charged aspartate at position 279 is less favorable than the smaller alanine, suggesting that DZ-modulation is inhibited by side chain interactions at the center of the linker. Fourth, α_1_V279D severely impairs unliganded gating. Fifth, α_1_V279A similarly enhances DZ-potentiation of GABA-evoked currents in a wild-type background. Sixth, the effects of α_1_V279A in both α_1_L9’T and wild-type backgrounds are explained by specific changes in the pore gating equilibrium and its coupling to DZ binding at the BZD site.

The use of gain of function mutations to study modulatory or weakly activating ligands is well appreciated (Akk et al., 2018; Campo-Soria et al., 2006; Downing et al., 2009; Rüsch and Forman, 2005; Findlay et al., 2001). Single channel gating dynamics for combinations of gain of function mutations in AChR suggest that the unliganded and agonist-bound gating mechanisms are similar if not identical, and also that they are similarly affected by mutations (Purohit and Auerbach, 2009). Consistent with this idea, MWC models of pseudo steady-state GABA_A_R function have been largely successful in describing the effects of gain of function mutations with changes to the pore gating equilibrium independent of agonist binding (Steinbach and Akk, 2019; Downing et al., 2009; Campo-Soria et al., 2006; Rüsch and Forman, 2005; Chang and Weiss, 1999). The model reported here in Figure 9 supports this conclusion.

Although the effects of α_1_V279A can also be explained by specific changes to the pore gating equilibrium, they are dependent on DZ occupation of the BZD site. From the model in Figure 9A we compute an efficiency for GABA of *η*_*GABA*_ = (1 − *log*(*K*_*G*_)/*log*(*cK*_*G*_)) × 100% = 37%, similar to that reported previously (Nayak et al., 2019). In contrast, DZ efficiency is *η*_*DZ*_ = (1 − *log*(*K*_*D*_)/*log*(*dK*_*D*_)) × 100% = 3%. Whereas α_1_V279A decreases unliganded pore opening, it also increases DZ efficiency (*η*_*DZ,V*279*A*_ = 9%) to the extent that DZ-binding overcomes its intrinsic inhibitory effect and results in similar or even enhanced activation as compared to wild-type (Figure 9). Interestingly, an M2-M3 linker mutation in the γ_2_ subunit was similarly observed to impair activation by GABA and enhance modulation by the general anesthetic propofol (O’Shea et al., 2009).

Our estimates for the energetic effect of DZ binding on pore gating (0.7 *kT* ≈ 0.4 *kcal/mol*) are similar to prior estimates from direct gating in L9’S/T backgrounds and from kinetic models (Goldschen-Ohm et al., 2014; Gielen et al., 2012; Downing et al., 2009; Rüsch and Forman, 2005). This is approximately 10-fold less than the energy derived from binding each molecule of GABA (Goldschen-Ohm et al., 2014; Maksay, 1994; Jackson, 1992). However, it is sufficient to produce a relevant change in open probability in channels with small free energy differences between closed and open states, such as conferred by single bound agonists, partial agonists, other allosteric modulators or gain of function mutations. From a clinical perspective, such small perturbations are likely to be more easily tolerated in patients. Given that the M2-M3 linker is associated with early movements during AChR activation (Purohit et al., 2013), it is possible that biasing the linker towards its active conformation in the α_1_L9’T background could limit our ability to observe its full range of motion. In this case our observed effects of linker mutations on DZ-modulation may underestimate their effects on the full activation pathway.

The interface between the extracellular agonist- and BZD-binding domains and the transmembrane helices is formed largely by several loops including the M2-M3 linker, Cys loop, β1-2 loop and loop F from neighboring subunits (Figure 1). Mutagenesis suggests that interactions between these domains are important for channel gating by agonists (Kash et al., 2003; O’Shea and Harrison, 2000) thought to involve an outward radial displacement of the M2-M3 linker (Nemecz et al., 2016). Rate versus free energy linear relationships in AChR suggest that the M2-M3 linker moves early during channel activation, similar to rearrangements at agonist binding sites (Purohit et al., 2013). Consistent with these observations, mutant cycle analysis indicates strong long-range coupling between residues in the M2-M3 linker and agonist binding sites (Gupta et al., 2017). Given the homology between the BZD site at the α/γ interface and agonist sites at β/α interfaces, interactions between the M2-M3 linker and BZD site are reasonable, although they need not be very strong given the relatively small energy DZ contributes to pore gating. As with agonist sites, such interactions could be mediated by global backbone conformational fluctuations or via distinct structural components, or both. Either way, side chain interactions at position 279 in the α_1_ subunit play an important role in coupling between the BZD site and pore gate.

In structural models the M2-M3 linker adopts a C-shaped conformation with α_1_V279 oriented inwards towards the center of the arc (Figure 1D) (Kim et al., 2020; Laverty et al., 2019; Masiulis et al., 2019; Phulera et al., 2018; Zhu et al., 2018). We hypothesize that the side chain at position 279 is centrally involved in steric interactions between linker residues near the top of the M2 and M3 helices. Removing this obstruction (α_1_V279A) would allow the linker to become more compact and bring the M2 and M3 helices closer together. In contrast, similar or larger side chains (α_1_V279D/W) would sterically inhibit such a conformational change. In this case, we speculate that the closer proximity of the M3 helix could impair unliganded pore gating by hindering radial expansion of the pore lining M2 helix. Conversely, a more compact transmembrane domain may also enhance coupling with the BZD site (Kim et al., 2020).

Alternatively, removing most of the central side chain may simply alter linker flexibility. Linker flexibility has been suggested to be inversely correlated with gating, where stabilizing interactions between α_1_R19’ (located just below α_1_V279) and the linker backbone may promote coupling between extracellular and intracellular domains (Masiulis et al., 2019). Thus, the strong inhibition of unliganded opening conferred by α_1_V279D could reflect competition for α_1_R19’, thereby weakening its interaction with the linker backbone and increasing linker flexibility. However, this does not provide a simple explanation for the opposing effects of α_1_V279A on unliganded versus DZ-bound gating. Regardless, it is important to keep in mind that the energetic changes that we observe for DZ-gating are on the order of a hydrogen bond or two, and thus can be accounted for by relatively subtle changes.

Loop F and pre-M1 in the neighboring γ_2_ subunit come in close proximity to the α_1_M2-M3 linker and are known to be important for BZD modulation, with loop F also contributing to the BZD binding site (Hanson and Czajkowski, 2011, 2008). In AChR strong coupling to the adjacent loop F and pre-M1 segment of the neighboring subunit occurs primarily via the threonine aligning with α_1_V279 in GABA_A_R and its neighboring serine (Gupta et al., 2017). A reorientation of the linker due to removal of steric interactions near its center could alter coupling to the BZD site via interactions with γ_2_ loop F and/or pre-M1. For example, a compression of the ends of the M2-M3 linker would cause the middle of the linker to be squeezed outward towards the neighboring γ_2_ subunit, potentially enhancing intersubunit coupling. However, other regions including the β4-5 linker in the channel’s outer vestibule also affect BZD modulation of agonist-evoked currents (Pflanz et al., 2018; Venkatachalan and Czajkowski, 2012). Thus, BZD modulation likely involves larger domain fluctuations in addition to any specific molecular pathways involving α_1_V279.

Recent structures of αβγ receptors show DZ bound not only at the classical α/γ site in the extracellular domain, but also in the transmembrane domain below the M2-M3 linker at β/α and γ/β intersubunit interfaces (Kim et al., 2020; Masiulis et al., 2019). Thus, it is intriguing to speculate that enhanced DZ-gating as conferred by α_1_V279A may involve occupation of a transmembrane site. However, these structures were solved in the presence of 100-200 μM DZ, and the relevance of these transmembrane sites to high affinity binding of ∼1 μM DZ at the classical site is unclear (Walters et al., 2000). Also, DZ binding in the transmembrane domain was not observed at α/γ or α/β interfaces. Therefore, we favor the interpretation that our observed effects reflect altered functional coupling with the classical site.

Our observations reveal a critical residue V279 in the α_1_M2-M3 linker which regulates energetic coupling between the BZD site and the pore gate, whereas other linker side chains contribute little or not at all. These data shed new light on the molecular basis for GABA_A_R modulation by one of the most widely prescribed classes of psychotropic drugs.

## Methods

### Mutagenesis and Expression in Oocytes

DNA for wild-type GABA_A_R rat α_1_, β_2_ and γ_2L_ subunits were a gift from Dr. Cynthia Czajkowski. Single alanine substitutions were introduced throughout the α_1_M2-M3 linker from L276-T283 in addition to the gain of function α_1_L9’T pore mutation (rat α_1_L263T) (QuikChange II, Qiagen). Each construct was verified by forward and reverse sequencing of the entire gene. mRNA for each construct was generated (mMessage mMachine T7, Ambion) for expression in Xenopus laevis oocytes (EcoCyte Bioscience, Austin, TX). Oocytes were injected with 27-54 ng of total mRNA for α, β and γ subunits in a 1:1:10 ratio (Boileau et al., 2002) (Nanoject, Drummond Scientific). Oocytes were incubated in ND96 (in mM: 96 NaCl, 2 KCl, 1 MgCl2, 1.8 CaCl2, 5 HEPES, pH 7.2) with 100 mg/ml gentamicin at 18°C.

### Two-Electrode Voltage Clamp Recording and Analysis

Currents from expressed channels 1-3 days post injection were recorded in two-electrode voltage clamp (Dagan TEV-200, HEKA ITC and Patchmaster software). Oocytes were held at -80 mV and perfused continuously with buffer (ND96) or buffer containing PTX, GABA or DZ. PTX was diluted from a 1M stock solution in DMSO. DZ was diluted from a 10 mM stock solution in DMSO. Fresh PTX and DZ stock solutions were tested several times with no change in results. GABA was dissolved directly from powder. A microfluidic pump (Elveflow OB1 MK3+) and rotary valve (Elveflow MUX Distributor) provided consistent and repeatable perfusion and solution exchange across experiments, which limited solution exchange variability to primarily differences between oocytes only. 10 second pulses of PTX, GABA or DZ were followed by 3-6 minutes in buffer to allow currents to return to baseline. Recorded currents were analyzed with custom scripts in MATLAB (Mathworks). Recordings of concentration-response relations were bookended by pulses of PTX to correct for any drift or rundown during the experiment. Briefly, currents were baseline subtracted to correct for drift and then scaled by a linear fit to the peak of each PTX response to correct for rundown (Supplementary Figure 2-1). Concentration-response relations were fit to the Hill equation:

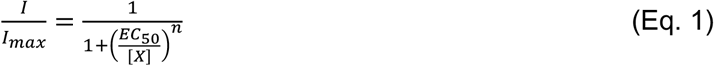

where *I* is the peak current response, [*X*] is ligand concentration, *EC*_50_ is the concentration eliciting a half-maximal response, and *n* is the Hill slope.

### DZ-Gating Model

For the model in Figure 7A, the free energy difference for unliganded (ΔG_0_) and DZ-bound (ΔG_1_) gating was estimated as:

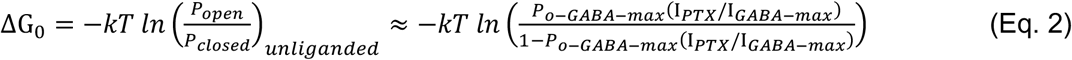

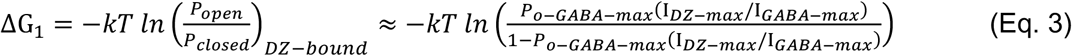

where I_*PTX*_, I_*GABA*_, and I_*DZ*_ are as shown in Figures 3 and 5, k is the Boltzmann constant, T is temperature, and *P*_*o*−*GABA*−*max*_ is the open probability in saturating GABA, which was assumed to be approximately 1. Importantly, even if this assumption is incorrect, the effect on DZ’s energetic contribution to pore gating should be minimal and our overall conclusion unchanged (see Figure 6B).

### MWC Model

For the model in Figure 9A, the probability to be in an open (O) state is given by:

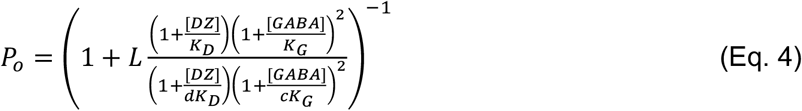

where L is the ratio of closed (C) to open (O) state probabilities in the absence of ligand, K_G_ and K_D_ are the respective dissociation constants for GABA or DZ, and c and d are the respective factors by which GABA or DZ binding influence pore opening.

## Acknowledgements

We thank Dr. Cynthia Czajkowski for gifting DNA for wild-type GABA_A_R subunits, and Drs. Richard Aldrich and Eric Senning for helpful discussions and shared use of equipment.

## Author Contributions

J.W.N carried out the experiments, analyzed the data and contributed to writing the manuscript. S.G. performed all molecular biology including generating DNA and RNA for each construct. M.P.G conceived and designed the project and contributed to data analysis and writing of the manuscript.

---

**Supplementary Figure 2-1.**
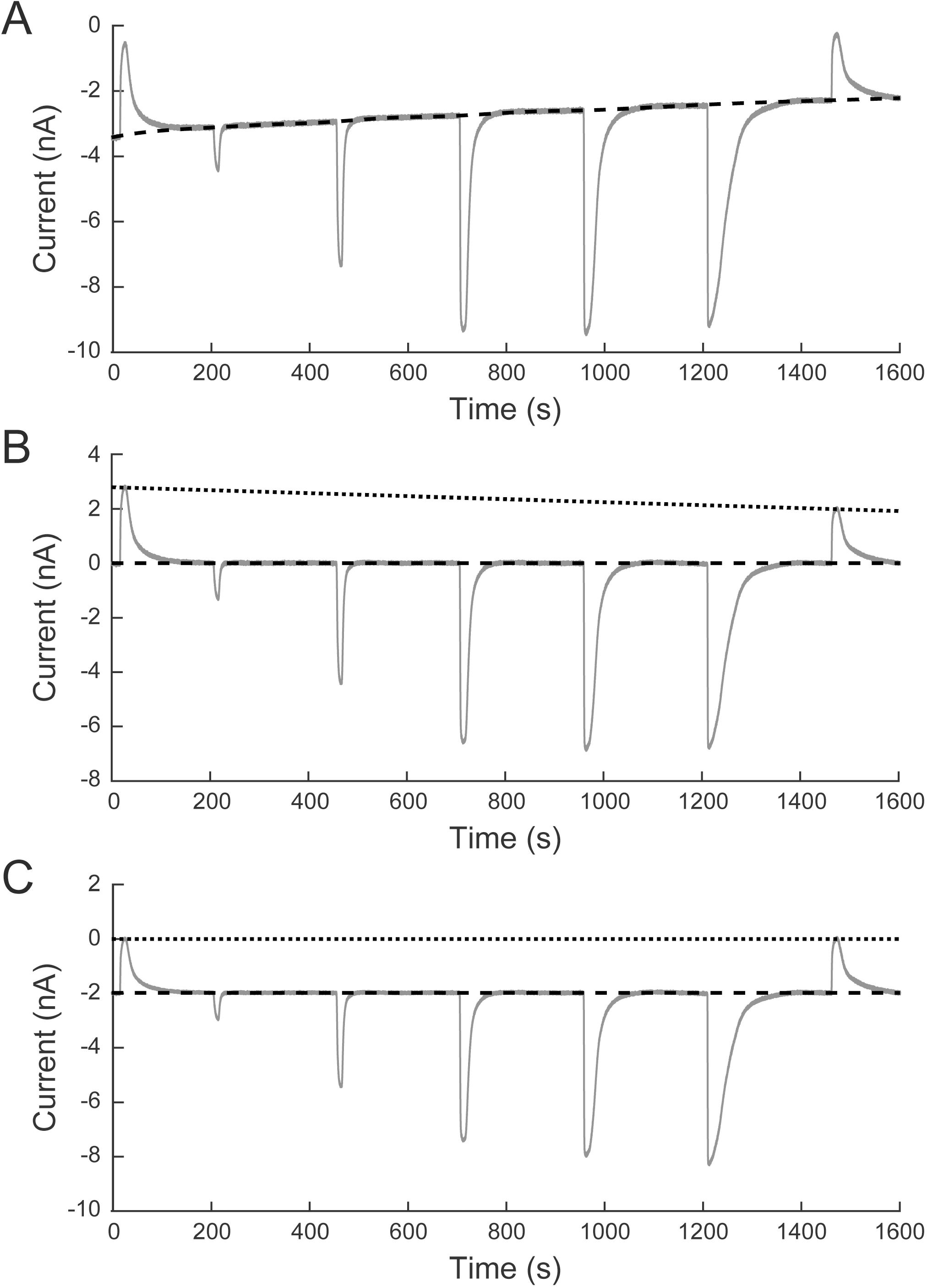
Correction for baseline drift and current rundown. **A)** Raw two-electrode voltage clamp current recording from α_1_L9’T/K278Aβ_2_γ_2L_ receptors expressed in a Xenopus laevis oocyte. Responses to 10 second pulses of increasing concentrations of GABA (0.1, 1, 10, 100 and 1000 μM) were bookended by 10 second pulses of 1 mM PTX as shown in Figure 2. **B)** Baseline drift was corrected by subtracting a spline fit to selected segments of baseline (dashed line in *A*). **C)** Current rundown was corrected by linearly scaling the current record so that the bookending PTX responses were of equal amplitude (dotted line in *B*). The resulting current trace was shifted so that zero current coincides with the peak response to block by PTX.

**Supplementary Figure 2-2.**
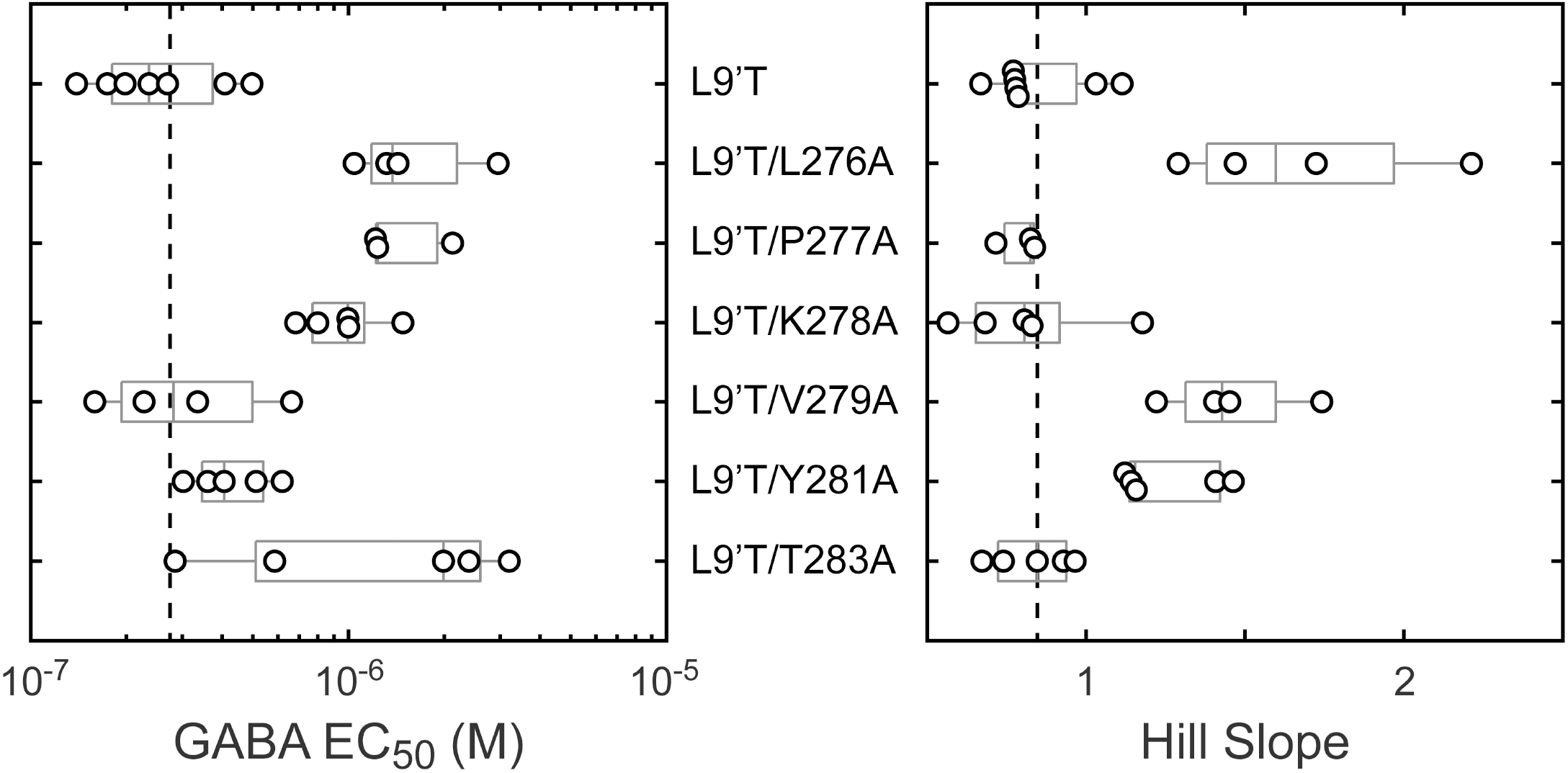
Summary of GABA EC_50_ and hill slope from fitting Equation 1 to GABA concentration-response relations as shown in Figure 2 for individual oocytes. Gray box plots indicate the median and 25^th^ and 75^th^ percentiles. The vertical dashed line is the mean for L9’T.

**Supplementary Figure 4-1.**
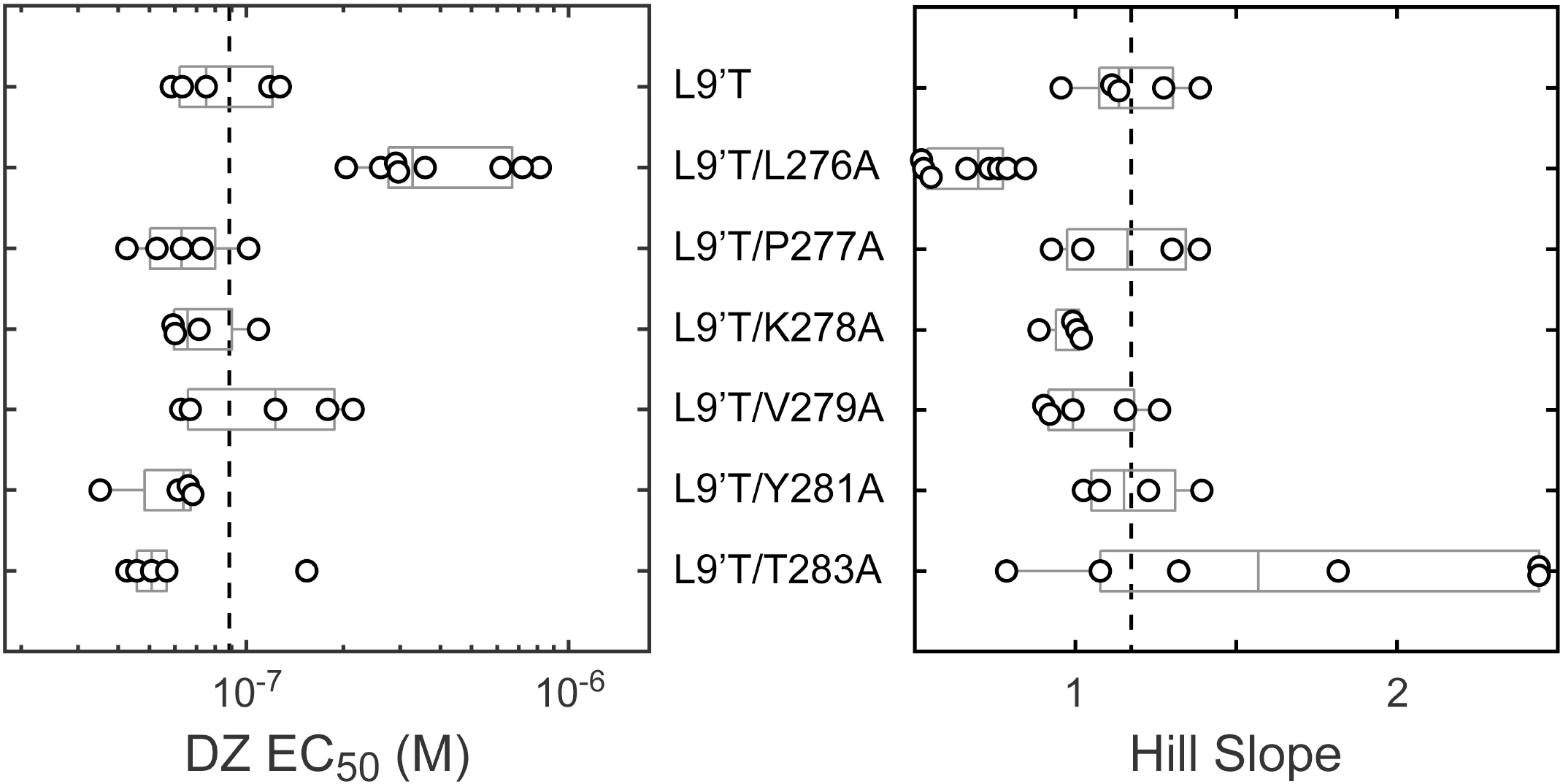
Summary of DZ EC_50_ and hill slope from fitting Equation 1 to DZ concentration-response relations as shown in Figure 4 for individual oocytes. Gray box plots indicate the median and 25^th^ and 75^th^ percentiles. The vertical dashed line is the mean for L9’T.

